# Optimizing in vitro spherulation cues in the fungal pathogen *Coccidioides*

**DOI:** 10.1101/2024.06.06.597856

**Authors:** Christina Homer, Elena Ochoa, Mark Voorhies, Anita Sil

## Abstract

*Coccidioides spp*. are part of a group of thermally dimorphic fungal pathogens, which grow as filamentous cells (hyphae) in the soil and transform to a different morphology upon inhalation into the host. The *Coccidioides* host form, the spherule, is unique and highly under characterized due to both technical and biocontainment challenges. Each spherule arises from an environmental spore (arthroconidium), matures, and develops hundreds of internal endospores, which are released from the spherule upon rupture. Each endospore can then go on to form another spherule in a cycle called spherulation. One of the foremost technical challenges has been reliably growing spherules in culture without the formation of contaminating hyphae, and consistently inducing endospore release from spherules. Here, we present optimization of in vitro spherule growth and endospore release, by closely controlling starting cell density in the culture, using freshly-harvested arthroconidia, and decreasing the concentration of multiple salts in spherulation media. We developed a minimal media to test spherule growth on various carbon and nitrogen sources. We defined a critical role for the dispersant Tamol in both early spherule formation and prevention of the accumulation of a visible film around spherules. Finally, we examined how the conditions under which arthroconidia are generated influence their transcriptome and subsequent development into spherules, demonstrating that this is an important variable to control when designing spherulation experiments. Together, our data reveal multiple strategies to optimize in vitro spherulation growth, enabling characterization of this virulence-relevant morphology.

## Introduction

*Coccidioides spp*. are dimorphic fungal pathogens found in the soil in desert regions in North, Central and South America^1^. In the soil, they grow as hyphae and, upon inhalation by a mammalian host, form a unique host-associated morphology known as the spherule^2^. Mature spherules are filled with hundreds of endospores and release these internal cells when they rupture. Each endospore can go on to form another spherule in a cycle called spherulation. *Coccidioides* causes infection in immunocompetent and immunocompromised individuals^3^, but efforts to develop new treatments and prevention strategies have been hindered by a lack of molecular knowledge of the spherule. This host form of the fungus has been difficult to study due in part to limited in vitro culture techniques. Groundbreaking early work by a number of groups established in vitro spherulation culture in Converse media^4-7^, RPMI tissue-culture media^8^, and co-culture with HeLa cells^9^. Since then, Converse media has become widely used but the components of this defined media that are essential for spherulation have not been further identified, and a more minimal media has not been established. Additionally, since microscopy techniques were limited and quantification was not common, many early reports did not quantify number of hyphae in spherulation cultures but relied on qualitative metrics. Two reports of continuous pure spherule culture (without hyphae) for 84 days^8^ and 4 years^10^ were achieved using initially laborious repeated filtration to remove hyphae, which would not be scalable to high-throughput applications. Thus, it remains challenging to grow a pure spherule culture that lacks hyphae. Finally, it is difficult to consistently induce endospore release without changing media or cell density^11^, thereby introducing additional variables into an experiment.

Here, we optimized in vitro spherulation, developed a minimal media for testing which carbon and nitrogen sources can be used for spherule growth, clarified a role for Tamol in dispersing a large visible film surrounding spherules, and developed a modified medium which consistently induces endospore release from spherules without nutrient supplementation or changing cell density. We additionally found that commonly used storage conditions greatly alter the transcriptome of arthroconidia. Finally, we found that the temperature used to generate arthroconidia affects both their viability and ability to form spherules. Together, this work represents an advance in the ability to consistently produce a pure spherule culture that reliably undergoes endospore release and increases our understanding of the cues required for spherule development.

## Methods Strains

The wildtype *Coccidioides posadasii* strain Silveira (NR-48944) was used for all experiments^12^.

### Arthroconidia Generation

Arthroconidia were inoculated onto 2x GYE agar (2 % Dextrose (Fisher), 1 % Yeast extract (Gibco), 1.5 % Bacto-Agar (BD)) + Penicillin/Streptomycin (100 U/ml penicillin and 100 g/ml streptomycin, UCSF Media Core) in T225 tissue culture flasks and grown for 4-6 weeks at 30°C, until the hyphal mat appeared dry and flattened as previously described^11^. Arthroconidia were harvested 0-2 days prior to initiating spherulation and stored at 4°C until use unless noted in the text. Arthroconidia harvest was done as previously described^11^, by adding PBS (UCSF Media Core) to tissue culture flasks with the hyphal mat, scraping to resuspend, and filtering through a 70-micron mesh filter. Arthroconidia were then washed twice with PBS and resuspended in PBS at appropriate concentrations for downstream assays. Arthroconidia were quantified using a plastic hemacytometer (Neubauer, C-Chip) sealed with nail polish.

### Microscopy of Spherulation and Hyphal Growth

Standard spherulation conditions: 125 mL polypropylene flasks containing 50 mL of Converse media inoculated with 10^6^/mL arthroconidia (unless otherwise stated) and placed at 39°C, 10 % CO_2_, shaking at 120 rpm. Converse media was made as previously published^11^ except for alteration in Tamol concentration (0.016 M NH_4_CH_3_CO_2_ (Sigma Aldrich), 0.0037 M KH_2_PO_4_ anhydrous (Fisher), 0.003 M K_2_HPO_4_ anhydrous (Fisher), 0.0016 M MgSO_4_*7H_2_O (Fisher), 1.25 × 10^−5^ M ZnSO_4_*7H_2_O (Fisher), 2.4 × 10^−4^ M NaCl (Fisher), 2.04 × 10^−5^ M CaCl_2_*2H_2_O (Fisher), 1.43 × 10^−4^ M NaHCO_3_ (Fisher), 0.5% Tamol^®^ (Northeast Laboratory), 0.4 % glucose (Fisher), 0.5 % N-Z amine (Fisher)). All stocks/media were sterile-filtered using a 0.45 micron vacuum filter instead of being autoclaved. Where noted, alterations were made to the Converse media including removing/reducing concentrations of individual components. Additional ingredients used in Converse alterations: 0.016 M KCH_3_CO_2_ (Sigma Aldrich), 0.016 M NaCH_3_CO_2_ (Fisher), 0.016 M NH_4_Cl (Fisher), 0.016 M (NH_4_)_2_SO_4_ (Molecular Sigma), 0.016 M NH_4_HCO_3_ (Sigma Aldrich). For hyphal growth, 125 mL polypropylene flasks containing 50 mL of Converse media were inoculated with 10^6^ arthroconidia/mL and grown at 25**°**C shaking at 120 rpm. At stated timepoints for light microscopy (Day 2-3, Day 4-5, and Day 7 post-inoculation), cells were fixed with 4 % paraformaldehyde (PFA) (Electron Microscopy Sciences) at room temperature for 30 minutes and washed twice in PBS, pelleting cells by centrifugation for 2 minutes at maximum speed between washes. Cells were visualized using a 40X DICII objective on a Zeiss Axiovert 200 microscope, with additional 1.6X Optovar magnification.

### Making endoConverse Media

Using Converse media as described above, pH was first raised to 11 with 4 N NaOH, then lowered to 8.5 with 6 N HCl. Visible precipitate was observed and then removed by sterile filtration with a 0.45 micron filter.

### Identifying Precipitate Produced During Generation of endoConverse Media

Using 100 mL of normal Converse, pH was increased to 11 with 4 N NaOH and then lowered to 8.5 with 6 N HCl. Precipitate was pelleted by centrifugation (maximum speed, 5 minutes) and the supernatant was removed. The precipitate was then sent to EAG Laboratories and analyzed by Scanning Electron Microscope/Energy Dispersive X-Ray Spectroscopy. Results presented in Figure 3C are Copyright © EAG, Inc. (www.eag.com).

### Arthroconidia Aging

For the experiment described in Figure 5, arthroconidia were harvested as described above and placed at 4°C in PBS (UCSF Media Core) in the dark. At specified timepoints, 3 aliquots were removed from the stock and RNA was immediately harvested as described below.

### Generating Arthroconidia at Different Temperatures

For the experiment described in Figure 6, arthroconidia were generated at either 25°C, 30°C, or 37°C for 4 weeks and then harvested as described above. 3 aliquots were removed from the arthroconidia stock and RNA was immediately harvested as described below. This experiment was repeated for a total of 2 replicates per stock generated at each temperature. One of the biological replicates harvested at 30°C is the same sample that was used for Week 0 in the arthroconidia aging experiment in Figure 5.

### RNA extraction and RNA-Seq Library Preparation

RNA from arthroconidia were collected from the same arthroconidia stock in triplicate by placing 5×10^7^ arthroconidia into Trizol LS (Ambion) and bead beating for 2 minutes. Samples were stored at -80°C until samples from all timepoints in an individual experiment had been collected. RNA was extracted using the Direct-zol RNA Miniprep Plus isolation kit (Zymo) with the on-column DNAse digestion step extended for 15 minutes. Sequencing libraries were prepared using the Ultra II Directional RNA Library Prep kit with dual-indexed multiplexing barcodes, with total RNA as the input, omitting the polyA selection step. Library quality and adaptor dimer contamination were analyzed using Agilent High Sensitivity DNA Chip. An additional round of library size selection was performed using homemade Serapure size selection beads^13^ for libraries containing significant adaptor dimers. Final library concentrations were measured using the Qubit High Sensitivity or Broad Range reagents. Libraries were pooled and sequencing was then performed on 3 and 2 lanes of the Novaseq 6000 S4 at the Chan Zuckerberg Biohub – San Francisco, for Figure 5 and Figure 6, respectively.

### RNA-seq data analysis

Analysis was conducted as previously described^14^ with alterations below. Briefly, estimated counts of each transcript were calculated for each sample by alignment-free comparison against the predicted mRNA for the published Silveira genome^12^ using KALLISTO version 0.46.2^15^.

Further analysis was restricted to transcripts with raw counts ≥ 10 in at least one sample across an individual experiment. Differentially expressed genes were identified by comparing replicate means for contrasts of interest using LIMMA version 3.46.0^16^. Genes were considered significantly differentially expressed if they were statistically significant (at 5 % false discovery rate) with an absolute log_2_ fold change ≥ 1 for a given contrast unless otherwise noted in the text. The Euler diagram in Figure 6 was generated using the eulerAPE package version 3.0.0^17^.

### Measurement of arthroconidia viability

Arthroconidia viability was measured using two different methods: 1) arthroconidia stocks were diluted 1:100 in 0.4 % Trypan Blue (Sigma) and counted on a hemacytometer. Arthroconidia were classified as viable if they excluded the blue dye and were classified as non-viable if they were blue. 2) Dilutions of arthroconidia stocks were plated on 2x GYE agar + Penicillin/Streptomycin, incubated at 30°C for 72 hours, and colonies were counted. Percent viability was defined as the number of colonies that had grown out of total number of arthroconidia plated (by hemacytometer counts).

## Results

### In vitro spherule formation depends on arthroconidia concentration

To study spherule morphology in detail, our first goal was to reproducibly generate spherules without hyphae also developing in the same culture. Arthroconidia germinate and develop into spherules, which go on to release endospores (Figure 1A). We tested spherule formation by placing arthroconidia in media known to induce spherules (RPMI + 10 % fetal bovine serum (FBS)^8,11^ or Converse media^4-7^) and germinated them in standard spherule-inducing conditions (39°C, 10 % CO_2_). We added 0.5 % Tamol to Converse media to encourage optimal growth^4^ (referred to as Converse media hereafter). Additionally, we placed arthroconidia in media similar to those used for macrophage infections with bone-marrow-derived macrophages (DMEM + 20% FBS^18-20^) and germinated them in spherule-inducing conditions. While spherules formed and released endospores in all three conditions (Figure S1), arthroconidia placed in Converse media formed the most robust spherules with very few hyphae so we chose this as our base media for further study. Next, given precedent that cell density can influence spherule formation and purity of spherule cultures^8^, we assayed whether cell density impacted the rate of spherule formation, the amount of hyphae in spherule cultures, and endospore release. We placed freshly harvested arthroconidia of varying densities in Converse media, germinated them in standard spherule-inducing conditions, and monitored spherule formation by light microscopy (Figure 1B). Lower density spherulation cultures were comprised of spherules, the vast majority of which went on to release endospores (quantification schema in Figure 1C, quantification data Figure 1D). Higher density cultures formed spherules, but those spherules did not go on to release endospores during the 7 days cultures were observed, perhaps due to nutrient depletion, accumulation of a metabolite that inhibits endospore release, or cell-density dependent signaling. We observed more hyphae in lower density cultures than higher density cultures. It is not clear whether those hyphae originated from newly released endospores or from the original arthroconidia, although hyphae generally appeared after endospore release. Cultures started at a concentration of 10^6^ arthroconidia/mL in Converse media and germinated under standard spherulation conditions released endospores starting on Day 3 and produced minimal numbers of hyphae. Therefore, we chose this as our standard arthroconidia concentration for starting spherule cultures.

**Figure 1:**
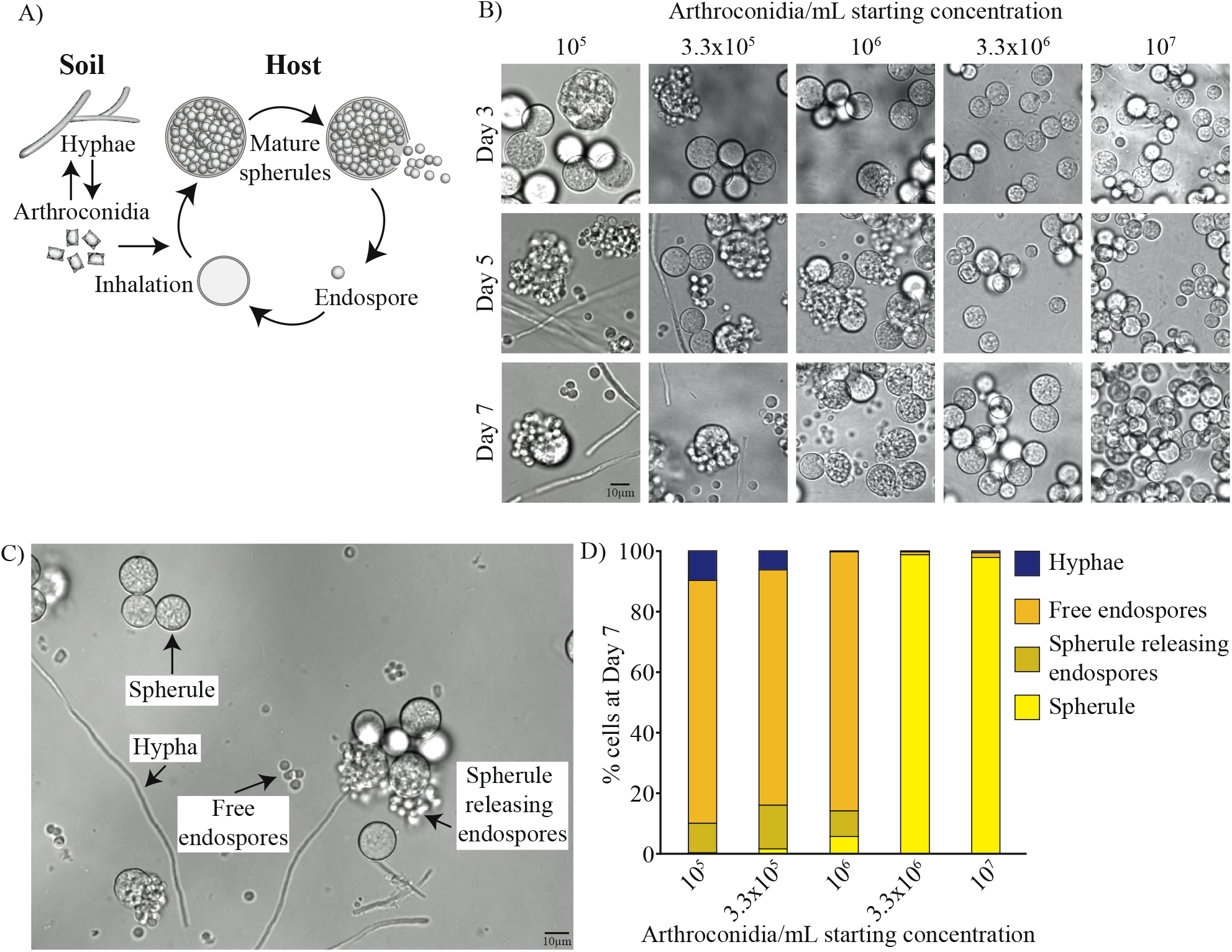
Optimizing starting concentration of arthroconidia for in vitro spherule formation. A. The schematic depicts the *Coccidioides* lifecycle. Hyphae grow in the soil and produce infectious spores named arthroconidia. If inhaled, arthroconidia enlarge into spherules that partition internally, generating endospores that are released when the spherule ruptures. Released endospores develop into new spherules and continue the cycle of spherulation. B. Different concentrations of arthroconidia ranging from 10^5^ – 10^7^ arthroconidia/mL were placed in Converse media + 0.5 % Tamol. Cells were fixed with 4 % PFA and monitored by light microscopy at day 3, 5, and 7 post-inoculation and monitored for spherule formation, hyphae formation, and endospore release. At higher densities, hyphae formation was suppressed but spherules did not release endospores during the experiment. At lower concentrations, there was more hyphal contamination of cultures but spherules did release endospores. C. DIC microscopy image of a spherulation culture fixed in 4 % PFA demonstrating all 4 possible morphologies for quantification schema: spherule prior to endospore release, spherule in the process of releasing endospores which are still associated with the spherule form, free endospores that have disassociated from the spherule that released them, and hyphae. D. Quantification of the proportion of each morphologic form in cultures on day 7 from B, using the schema demonstrated in C. Each morphology was quantified by hand for at least 10 fields of view for each sample.

### Developing a minimal media for spherulation

We wanted to create a minimal media for spherulation that could be used to test the effect of various carbon sources. Converse has 4 ingredients that are standard carbon sources: glucose, amines, ammonium acetate, and sodium bicarbonate. The chemical Tamol^4^ also contains carbon although, to our knowledge, it is not known whether organisms are able to use it as a carbon source. We placed arthroconidia in one of the following (Figure 2A, 2B): (1) Converse, (2) Converse media lacking glucose, amines, or ammonium acetate, (3) Converse lacking glucose and amines, or (4) Converse media lacking sodium bicarbonate. Spherules developed in all these media variations except for those lacking ammonium acetate, in which some cells appeared to be arrested at the stage of isotropic swelling or early spherule initials, in combination with ungerminated arthroconidia. Of note, spherule diameter in media lacking glucose was smaller than spherules in normal Converse or Converse lacking amines (Figure S2A). Since ammonium acetate could be promoting growth by serving as either a carbon source (acetate) or a nitrogen source (ammonium), we substituted other sources of acetate or ammonia for ammonium acetate and found that the growth defect in Converse lacking ammonium acetate could be partially or fully rescued by other chemicals containing ammonia (Figure 2C), indicating that ammonia appears to be the critical nitrogen source during spherulation despite the presence of amines in the media. As spherules grown in media containing ammonium chloride and ammonium sulfate instead of ammonium acetate are smaller than spherules in normal Converse, there is likely some role for acetate as a carbon source. Consistent with that, spherules grown in Converse containing ammonium bicarbonate instead of ammonium acetate grew to a similar size as spherules grown in Converse with ammonium acetate. This is likely because bicarbonate can also serve as a carbon source, like acetate.

**Figure 2:**
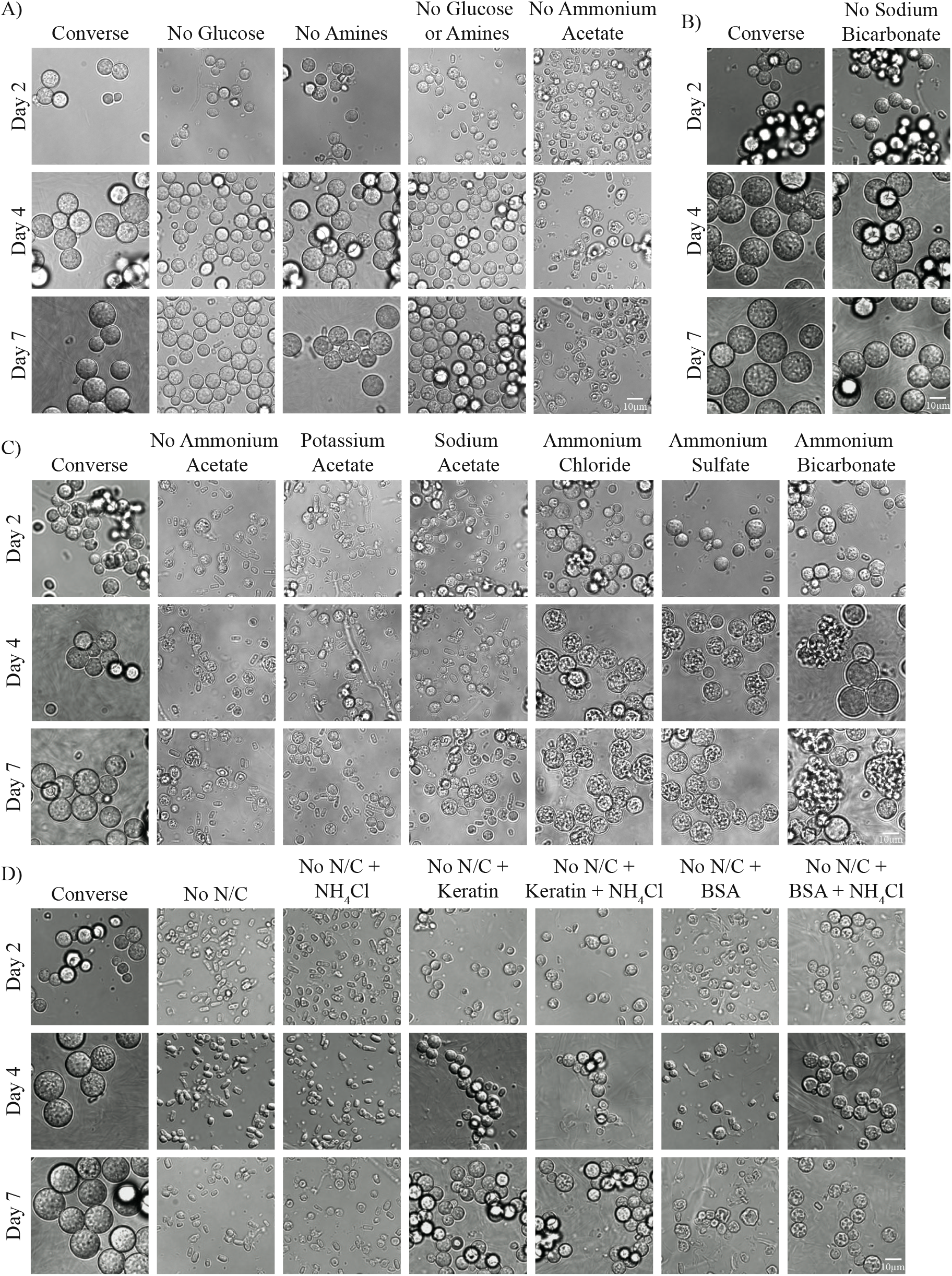
Defining carbon and nitrogen sources during spherulation. 10^6^ arthroconidia/mL were grown in standard spherulation conditions (39°C, 10 % CO_2_) in media variations described below and the resultant cells were fixed with 4 % PFA and monitored for spherule formation by light microscopy on day 2, 4, and 7 post-inoculation. A) Media was either standard Converse or Converse lacking glucose (No Glucose), lacking N-Z amines (No Amines), lacking both glucose and N-Z Amines (No Glucose or Amines), or lacking ammonium acetate (No Ammonium Acetate). B. Media was either standard Converse, or Converse lacking sodium bicarbonate. C. Media was either standard Converse, Converse lacking ammonium acetate (No Ammonium Acetate), or Converse in which the ammonium acetate is replaced by the same concentration of potassium acetate, sodium acetate, ammonium chloride, ammonium sulfate, or ammonium bicarbonate. D. Media was either Converse or No N/C (Converse media lacking carbon and nitrogen sources: glucose, N-Z amines, ammonium acetate, and sodium bicarbonate), supplemented with 3 % keratin, 3 % BSA, and/or 0.016 M NH_4_Cl and fixed/monitored as described in A. The biological replicate labeled as ‘Converse’ is the same biological replicate as presented in B.

**Figure 3:**
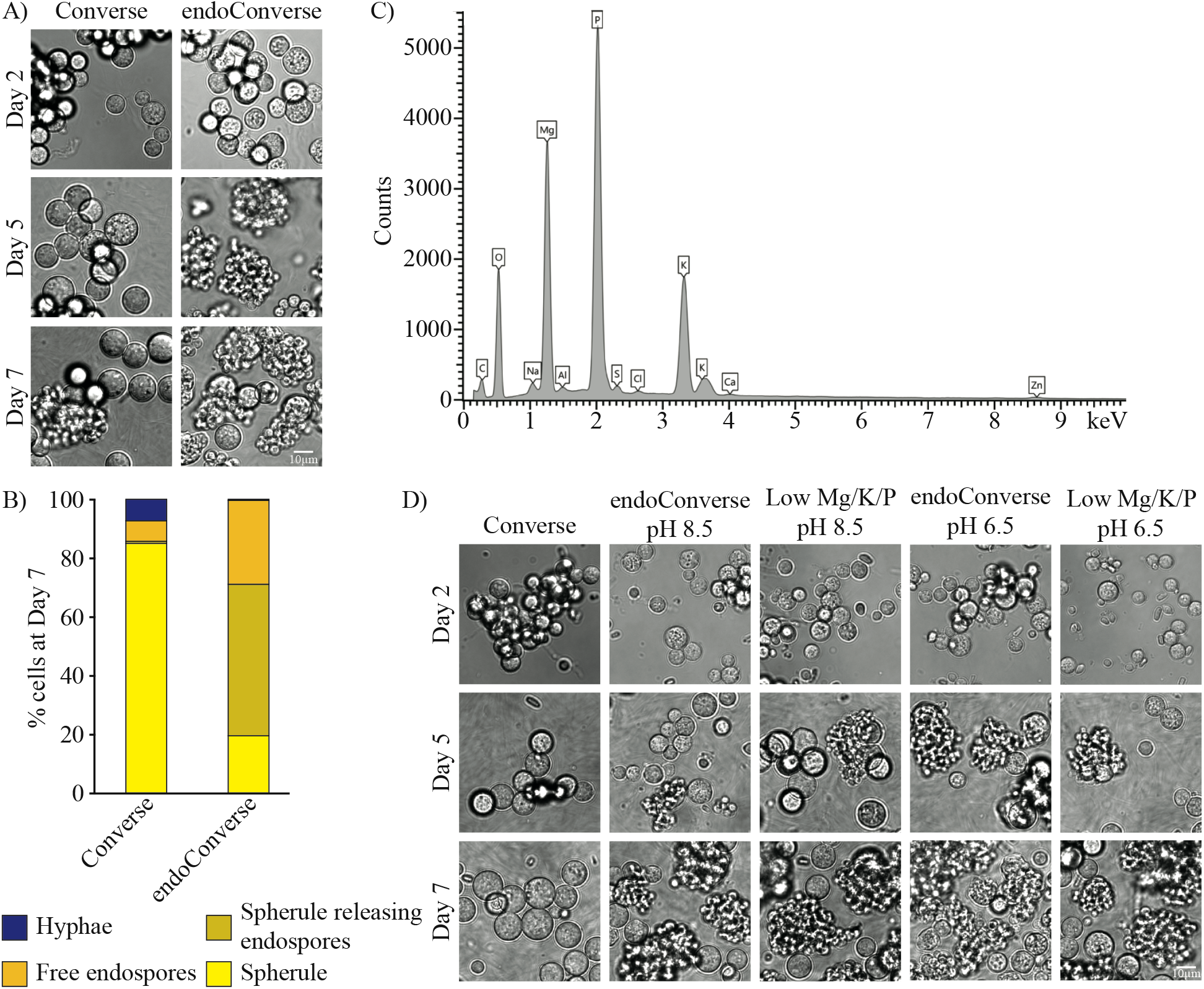
Optimizing endospore release. A. 10^6^ arthroconidia/mL were grown in standard spherulation conditions (39°C, 10% CO_2_) in either Converse or endoConverse. Cells were fixed with 4 % PFA and monitored by light microscopy at day 2, 5, and 7 post-inoculation for spherule formation. B. Quantification of the proportion of each morphology in cultures on day 7 from 3A (schema in 1C). Each morphology was quantified by hand for at least 5 fields of view for each sample. C. Identification of precipitate removed in the process of making endoConverse by energy dispersive X-ray spectroscopy. D. 10^6^ arthroconidia/mL were grown in standard spherulation conditions (39°C, 10 % CO_2_) in either Converse or endoConverse with starting pH 6.5 or 8.5, and Converse media with only 10 % the normal starting amount of MgSO_4_, KH_2_PO_4_ and K_2_HPO_4_. Cells were fixed by 4 % PFA and monitored by light microscopy at day 2, 5, and 7 post-inoculation for spherule formation. The biological replicate labeled as ‘Converse’ is the same biological replicate as presented in 2C.

Since our data indicate that spherules can use multiple carbon sources, we created a Converse media lacking glucose, amines, ammonium acetate, and sodium bicarbonate. As expected, arthroconidia placed into this media in spherulation conditions were not able to form spherules (and also do not form hyphae), indicating they were not able to germinate and grow (Figure 2D). Since we know ammonium acetate serves as an important nitrogen source as well, we added ammonium chloride to this media and found that it did not restore growth, suggesting the lack of growth is due to lack of carbon availability, despite Tamol being present in the media. Therefore, we conclude Tamol does not serve as a carbon source during spherulation. Finally, to demonstrate that this “no carbon” Converse can be used to test the efficacy of various carbon sources for spherulation, we added keratin or BSA to “no carbon” Converse with or without ammonium chloride to see if *Coccidioides* spherules are able to use these proteins as carbon sources. We observed small spherules forming in the media containing keratin with or without ammonium chloride supplementation, indicating that keratin can serve as both a carbon and a nitrogen source during spherulation (Figure 2D and S2B). However, in “no carbon” Converse supplemented with BSA, we only observed small misshapen forms, potentially spherule initials, that were not able to progress through development. When “no carbon” Converse with BSA was supplemented with additional nitrogen (ammonium chloride), small spherule initials were able to form although they had a different appearance with more accentuated septations than conditions containing keratin or normal Converse. The significance of this difference in appearance is not clear. Together, these data suggest *Coccidioides* spherules can use keratin as both a carbon and nitrogen source and potentially BSA as a suboptimal carbon source. The keratin findings are particularly intriguing given that 1) *Coccidioides* is enriched in the environment in rodent burrows^21,22^, which are abundant in keratin and 2) clinically, disseminated *Coccidioides* infections often involve the skin, which is a keratinous structure^23^.

### Optimizing in vitro endospore release

Endospore biology, including endospore release, has not been studied. To robustly study this part of the spherulation cycle, we wanted to develop in vitro conditions where endospore release occurs reliably. While many of our samples in Converse media in spherulation conditions release endospores between day 3 to 7, the degree of endospore release is variable and spherules releasing endospores can be a small percentage of the culture in some instances. We found that raising the pH of Converse to 11 causes a precipitate to form, and when that precipitate is removed through filtration and the pH adjusted to 8.5, the resulting media (which we call endoConverse) induces a dramatically increased amount of endospore release by day 5 (Figure 3A), with the vast majority of spherules releasing endospores on day 7 (Figure 3B). We identified the elemental components of this precipitate using energy dispersive X-ray spectroscopy. While this assay is not quantitative, it can distinguish major and minor components. The assay identified that the major components of the precipitate were phosphorus, magnesium, potassium, and oxygen (Figure 3C). Based on the ingredients present in Converse media, we hypothesize that the magnesium sulfate and potassium phosphate are combining to form magnesium potassium phosphate salt, which has been shown to precipitate at high pH^24^. We attempted to replicate this effective decrease in salts via precipitation by reducing the MgSO_4_, KH_2_PO_4_, and K_2_HPO_4_ to 10% of their amount in normal Converse and adjusted the pH of that media to 8.5 to match endoConverse. When arthroconidia were placed into this altered media and grown under standard spherulation conditions, they did have significantly increased endospore release compared to normal Converse (Figure 3B). In fact, this effect held when the starting pH was 6.5 as well, as in normal Converse. Further investigation is warranted into the endospore release signaling pathways that are inhibited by the Mg, K, P, O-containing salt, which may give insight into when and how endospore release is triggered during infection.

### Tamol dislodges a visible outer film from spherules

As Tamol is an unusual ingredient in growth media, we interrogated the role it plays in Converse media. First, we eliminated Tamol entirely from Converse and found that, while arthroconidia placed in spherulation conditions still formed small spherules, their growth was arrested compared to the same media with Tamol, and these small spherules did not go on to release endospores (Figure 4A). As Tamol was not found to be a carbon source or nitrogen source (Figure 2), we do not think Tamol is altering the size of spherules due to nutritional factors. Instead, we hypothesize that its characteristics as a dispersant are altering the spherule surface in some way that facilitates expansion of the spherule cell wall. Next, we queried a range of Tamol concentrations in Converse (from 0.05 % to 0.5 %) and found that even the lowest concentrations of Tamol were sufficient to form spherules that are a similar size to those grown in Converse with 0.5 % Tamol (Figure 4B), although only a very small amount of endospore release was observed in these flasks. However, at the two lowest concentrations of Tamol tested (0.05 % and 0.1 %) and in Converse lacking Tamol, we observed accumulation of a substance around multiple spherules (arrowheads). This film is larger and more pronounced in the lowest concentration of Tamol tested. To further evaluate this film, we induced spherulation in normal Converse or endoConverse and then shifted already-formed spherules on Day 3 into Converse without Tamol. In these cultures, clumps of spherules or clumps of spherules releasing endospores accumulated a large, highly visible layer of this substance around themselves (Figure 4C, Figure 4D). This is not an effect of the media exchange, as transferring spherules into Converse with 0.5 % Tamol on day 3 does not cause a similar effect (Figure 4E). While we have not characterized this film definitively, we hypothesize that it is the spherule outer wall (SOW), which has been previously identified as a structure that is shed into the media of spherulating cultures^25^, and that Tamol is increasing the rate of that shedding. Therefore, in cultures with low or no Tamol, the rate of SOW shedding is decreased to the point that it accumulates around spherules into a thick enough film to be visible by microscopy. Further work is necessary to confirm whether this accumulating layer is SOW and whether such a layer accumulates during infection in a mammalian host.

**Figure 4:**
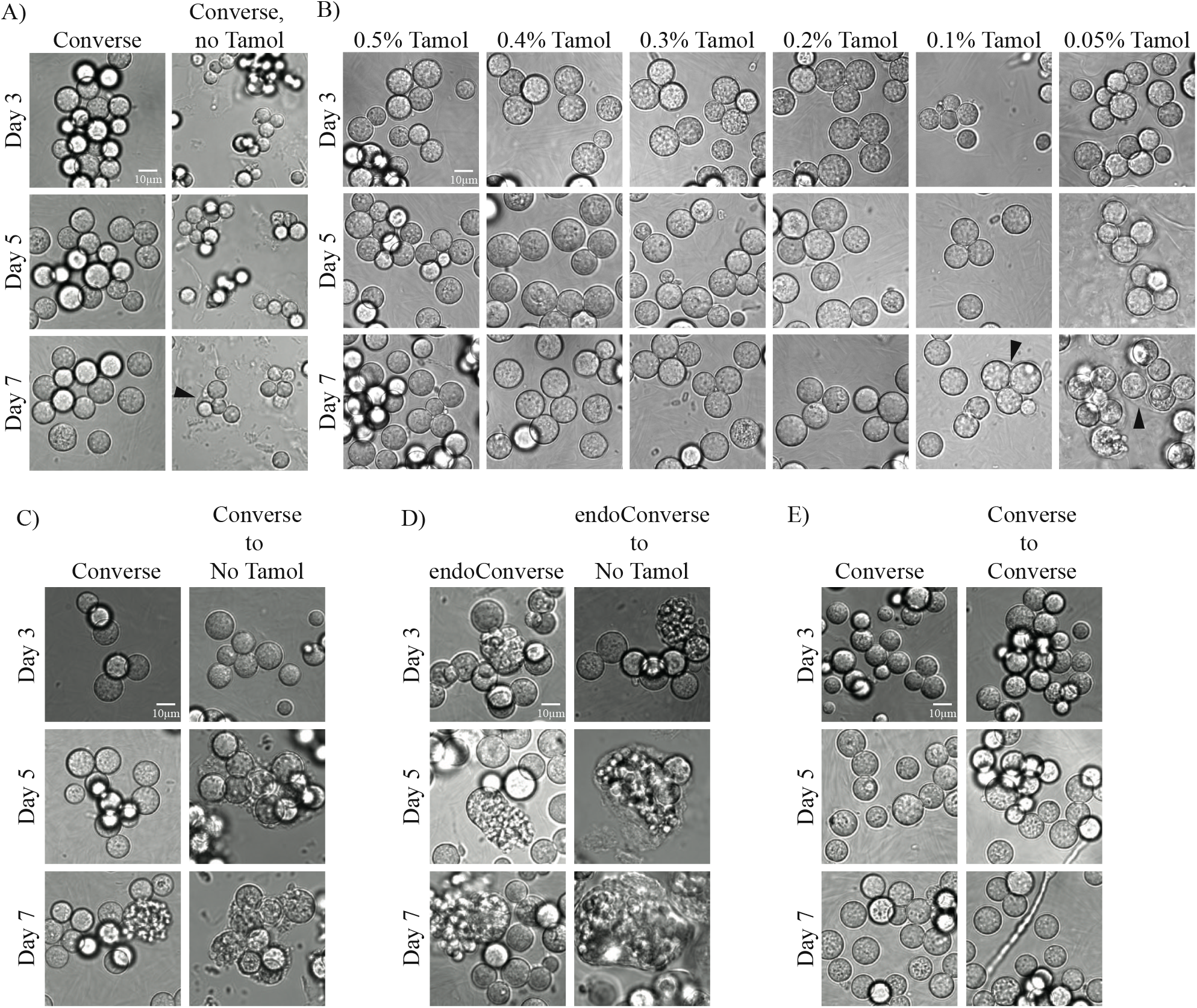
Dissecting the role of Tamol in spherulation. 10^6^ arthroconidia/mL were grown in standard spherulation conditions (39°C, 10 % CO_2_) in the media variations below. Cells were fixed with 4 % PFA and monitored by microscopy at day 3, 5, and 7 post-inoculation for spherule formation. A. Media was either Converse or Converse without Tamol. B. Media was either Converse or Converse with decreasing amounts of Tamol. The biological replicate labeled as ‘Converse’ is the same biological replicate as presented in 4A. Black arrowheads point to the layer observed to accumulate around groups of spherules in lowest concentrations of Tamol tested. C. Media was Converse at initiation of spherulation. On day 3, after cells were visualized by microscopy, samples labeled ‘Converse to No Tamol’ were spun down at 2500 rpm for 5 minutes, supernatant was discarded, and cells were resuspended in 50 mL of Converse lacking Tamol. They were then replaced at incubation conditions. D. Same as in 4C but spherules initiated with endoConverse instead of standard Converse media and exchanged to Converse lacking Tamol on day 3. E. Same as in 4C but exchanged to standard Converse on day 3.

**Figure 5:**
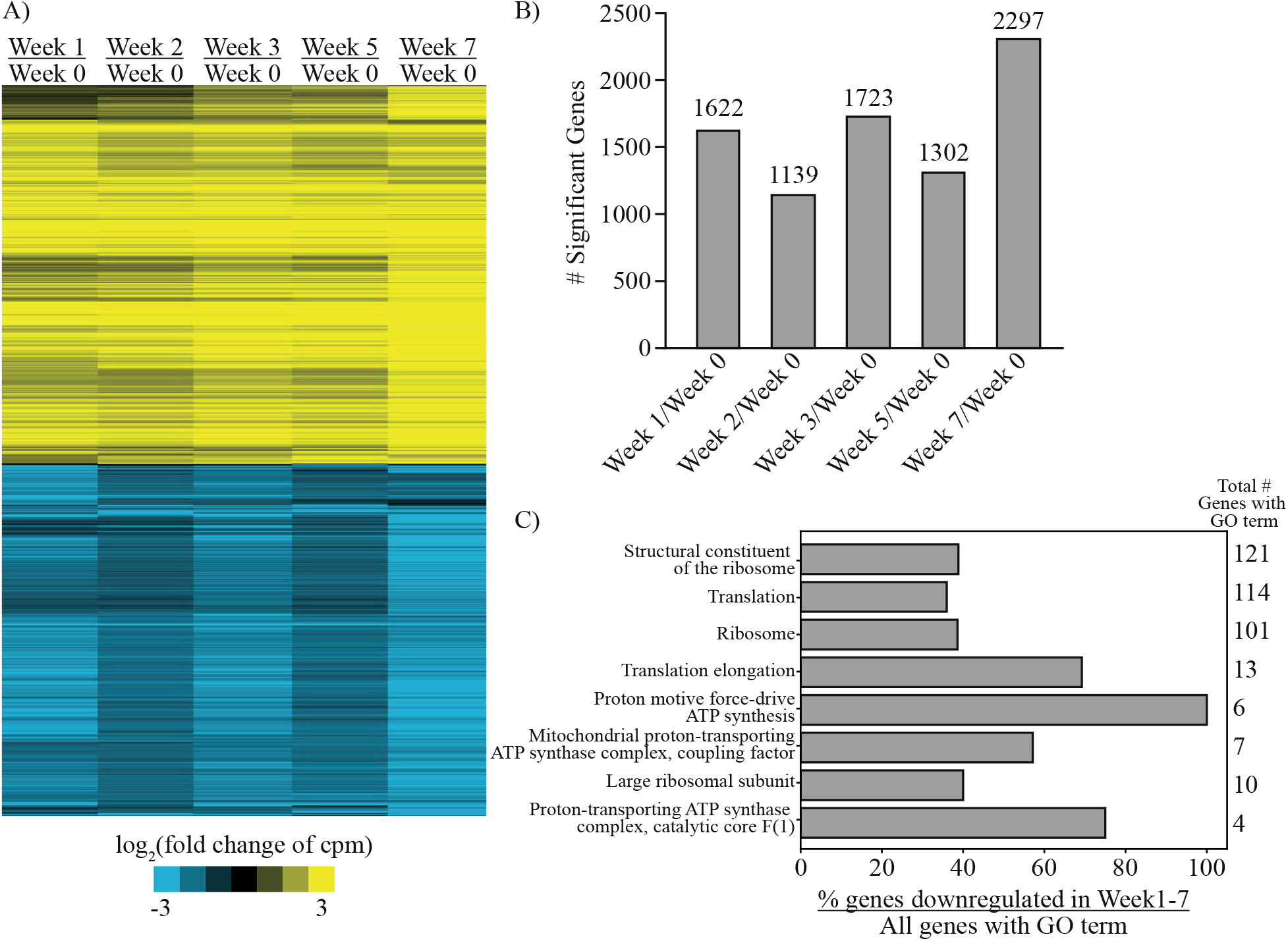
Storing arthroconidia in PBS at 4°C induces large-scale transcriptome changes. A. Heatmap demonstrating comparisons of the transcriptome between a stock of freshly harvested arthroconidia and arthroconidia stored at 4°C in PBS for 1, 2, 3, 5, and 7 weeks after harvest. Yellow (positive) and blue (negative) shades represent log_2_ fold change values of counts per million (cpm) for each stated comparison. Each row represents one transcript, rows were clustered based on behavior across all comparisons. B. Bar graph showing the number of significantly-differentially regulated transcripts (4-fold change, FDR 5 % in limma) for each comparison of triplicate samples at each stated timepoint. C. Percent of genes containing each GO term, with total genes in the genome containing that same GO term labeled on right of graph.

**Figure 6:**
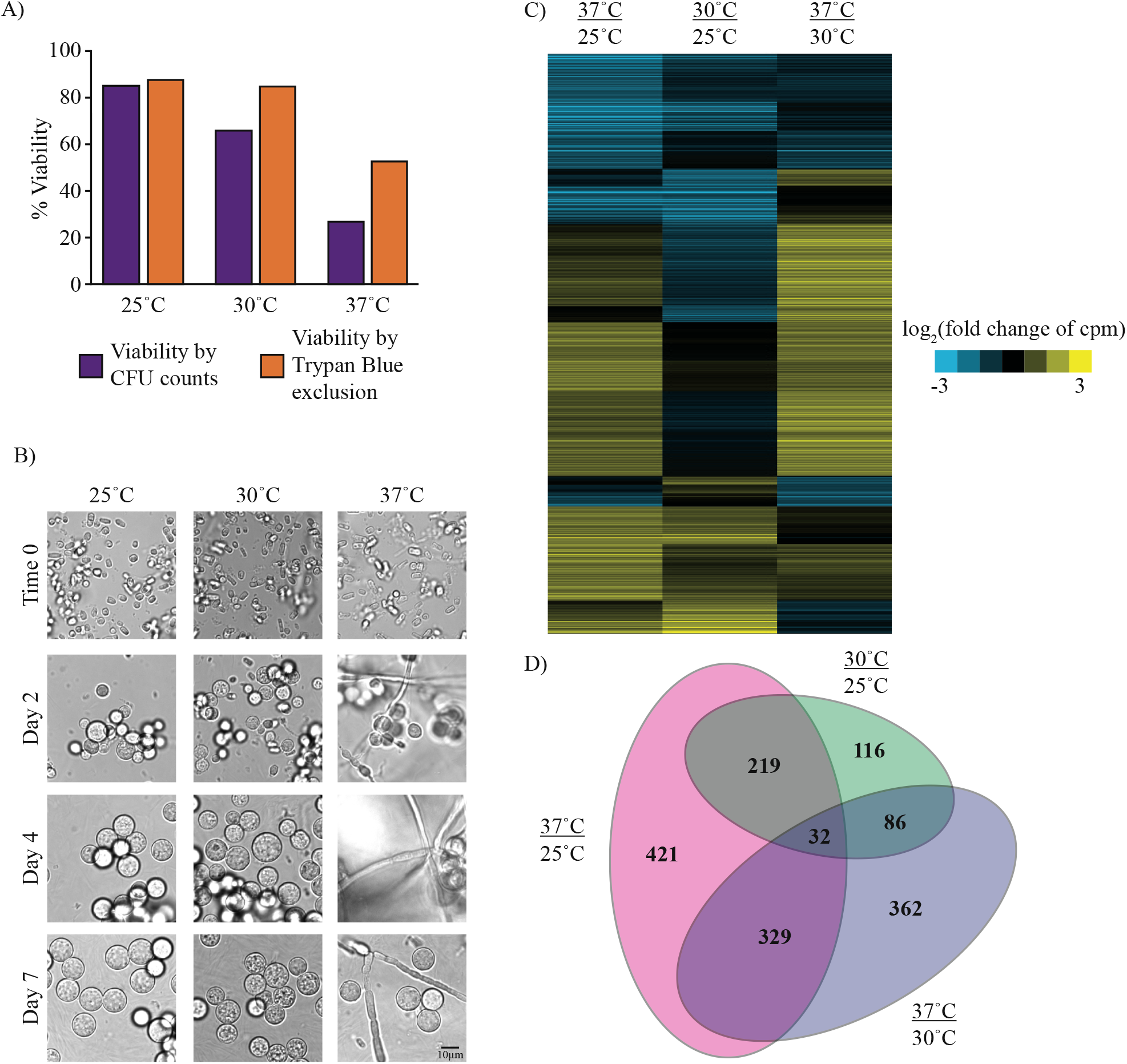
Generating arthroconidia at different temperatures affects their transcriptome and behavior upon germination. A. Percent of viable arthroconidia assessed by trypan blue exclusion (on day of arthroconidia harvest) or by CFU counts (after growth of arthroconidia stock for 72 h on 2x GYE at 30°C). B. 10^6^ viable arthroconidia/mL (as assessed by trypan blue), generated at either 25°C, 30°C, or 37°C, were grown in standard spherulation conditions (39°C, 10 % CO_2_) in Converse media. Cells were fixed with 4 % PFA and monitored by light microscopy at day 2, 4, and 7 post-inoculation for spherule and hyphae formation. C. Heatmap demonstrating comparisons of the transcriptome between arthroconidia stocks generated at 25°C, 30°C, and 37°C. Yellow (positive) and blue (negative) shades represent log_2_ fold change values of counts per million (cpm) for each stated comparison. Each row represents one transcript, rows were clustered based on behavior across all comparisons. D. Euler diagram demonstrating overlap between differential transcripts for comparisons between arthroconidia stocks generated at different temperature.

### Arthroconidia storage conditions greatly alter the transcriptome and affect in vitro spherule formation

The current standard method of storing *Coccidioides* arthroconidia stocks is in PBS at 4°C. We interrogated whether storage in these conditions alters the propensity of the arthroconidia stocks to form spherules when placed in our optimized spherulation conditions. Indeed, we found that the longer arthroconidia stocks were stored at 4°C in PBS, the more hyphae were present in spherule cultures generated from those stocks (Figure S3A). Given the precedent that the transcriptome of a fungal spore influences cellular phenotypes after spore germination^26,27^, we assayed the transcriptome of an arthroconidia stock after it was freshly harvested, or over time in storage. For stored arthroconidia stocks, we placed arthroconidia directly from PBS at 4°C into Trizol so that they did not have an opportunity to germinate. Indeed, we found that 2559 transcripts demonstrated a significant change in abundance between at least one timepoint in storage compared to freshly harvested arthroconidia (4-fold cutoff, FDR 5 %) (Figure 5A, Figure S3B, Table S1). There were similar numbers of transcripts whose abundance increases versus decreases during storage. Storage for 1-5 weeks at 4°C in PBS induced significant differences in transcript abundance for 1139 to 1723 transcripts (Figure 5B) and storage for 7 weeks induced the most change in transcript abundance (2297 transcripts). Overall, there was a similar trend in magnitude and direction of change for all transcripts responsive to placement in storage conditions.

While no clear gene ontology (GO) term enrichment was found in transcripts whose abundance was decreased upon storage, some transcripts related to sexual development and meiosis are, surprisingly, more abundant in 4°C storage conditions, including D8B26_004117 (ortholog of *Saccharomyces cerevisiae*’s *DMC1* meiotic recombinase^28^), D8B26_000913 (ortholog of *VELC*, a regulator of sexual development in *Aspergillus nidulans*^29^ *and Aspergillus fumigatus*^30^), D8B26_006129 (ortholog of *SPO11* in *S. cerevisiae*^31^, a meiosis-specific topoisomerase), D8B26_004631 (ortholog of *CSM3* in *S. cerevisiae*^32,33^,which plays a role in chromosome segregation during meiosis), D8B26_001535 (ortholog of *HOP1* in *S. cerevisiae*^34^, where it plays a role in homolog pairing during meiosis), and D8B26_04250 (ortholog of *MND1* in *S. cerevisiae*^*32,35*^, where it plays a role in recombination and meiotic nuclear division).

Examining the 264 transcripts that have significantly decreased abundance at all subsequent timepoints compared to Week 0, transcripts with GO terms related to the ribosome or translation are significantly enriched (Figure 5C). Given many of these downregulated transcripts are structural constituents of the ribosome, this likely implies a decrease in steady-state levels of ribosomes, but it remains to be explored whether this is due to a decrease in production of ribosomes or an increased rate of ribosomal transcript degradation. Almost all the genes containing GO terms related to ATP synthase are also downregulated upon storage. This may reflect a shift in energy sources or possibly a shift into dormancy, as is expected in a number of spore types^26,36,37^.

### Generating arthroconidia at different temperatures affects their transcriptome and behavior upon germination

Given that storage conditions affected the transcriptome of arthroconidia, we also sought to determine whether the conditions used to generate arthroconidia affected their transcriptome and behavior after germination. We generated arthroconidia by growing hyphal mats for 4 weeks at either 25°C, 30°C, or 37°C. After harvesting, we tested the viability of these arthroconidia stocks by trypan blue exclusion or quantifying the number of colony-forming units (CFUs) generated by each stock (Figure 6A, Figure S4A demonstrating additional replicate given variability in this assay). While viability measured by CFUs is consistently lower than by trypan blue, both methods indicate decreased viability of the arthroconidia stocks generated at 37°C compared to stocks generated at other temperatures. We then went on to characterize the ability of these arthroconidia stocks to generate spherules. Given our previous observations that lower concentrations of arthroconidia generate spherulation cultures containing more hyphae, we attempted to compensate for the decreased viability of arthroconidia generated at 37°C by inoculating with 10^6^ viable arthroconidia as assessed by trypan blue exclusion (since that measure of viability is available at the time of spherule culture initiation). Despite this compensation, spherulation cultures initiated with arthroconidia generated at 37°C had significantly more hyphae than arthroconidia stocks generated at 25°C and 30°C (Figure 6B). This may be because trypan blue staining overestimates arthroconidia viability (e.g. as compared to viability measurements by CFUs). However, the more intriguing possibility is that the conditions under which arthroconidia are generated are affecting spherulation behavior. Further study will be needed to verify this effect.

To delve deeper into the differences caused by generating arthroconidia at various temperatures, we assayed the transcriptome of these arthroconidia stocks, placing arthroconidia directly from the PBS in which they were harvested into Trizol so that they did not have the opportunity to germinate. We found that 1001 transcripts were significantly changed when comparing arthroconidia generated at 37°C versus 25°C. As expected, slightly fewer transcripts were significantly different when comparing the intermediate 30°C temperature: 809 transcripts were significantly differential between arthroconidia generated at 37°C versus 30°C, and 453 transcripts were significantly differential between arthroconidia generated at 30°C versus 25°C (2-fold cutoff, FDR 5 %, Figure 6C, 6D, Table S2). Indeed, the larger number of significantly differential transcripts in comparisons involving the 37°C-generated stock correlates with the more significant phenotypes of that arthroconidia stock (viability defect and hyphal contamination in spherule cultures). When focusing on the most drastic comparison of 37°C-generated arthroconidia compared to 25°C-generated arthroconidia, >50% of transcripts with the GO term 6270 (DNA replication initiation) are upregulated in the 37°C-generated arthroconidia, including D8B26_002886 (the ortholog of *CDC45* in *S. cerevisiae*, a DNA replication initiation factor^38^), D8B26_000131 (the ortholog of *SLD2*, which is required for chromosomal replication in *S. cerevisiae*^39^), D8B26_008289 (the ortholog of *CDC6*, a protein required for DNA replication initiation^40,41^ and that has been shown to interact with the eukaryotic replicative helicase in *S. cerevisiae*^42^) and multiple components of the eukaryotic replicative helicase (D8B26_000480, D8B26_006296, D8B26_004943). The significance of increased abundance of transcripts encoding replication factors in these presumably dormant arthroconidia is unknown and may suggest dysfunctional regulation of DNA synthesis in the 37°C-generated arthroconidia stocks that impacts their viability and potential to germinate into spherules.

## Discussion

Here we define conditions that result in optimal formation of endosporulating spherules. Optimizing in vitro spherulation and endospore release will pave the way for in-depth molecular studies of these infection-relevant morphologies. Current antifungals target conserved structures among fungi, but, given their lack of efficacy in treating coccidioidomycosis, pursuing a *Coccidioides*-specific target in the spherule or endospore may prove to be a more effective treatment. In addition, the minimal media developed here for in vitro growth of spherules will enable dissection of the carbon and nitrogen sources that are sufficient for spherule formation and will potentially identify those that are optimal for spherule formation as well. This information could provide insights into host niches that are particularly well-suited for spherule growth and persistence and could also lead to discovery of additional pathways that could serve as antifungal drug targets^43,44^. Given the observation that *Coccidioides* infection can be controlled for long periods of time while a patient is on antifungal treatment, but then the infection worsens or relapses upon stopping antifungal treatment^45^, it is hypothesized that there is a latent phase of *Coccidioides* infection^46-48^. Understanding this latent phase, and the alterations in metabolism associated with it, could hold the key to definitively curing disseminated infections^49^.

Our experiments also explored the role of Tamol in spherulation. Surprisingly, the removal of Tamol from spherulation media revealed a thick visible film surrounding spherules and endospores. This observation is reminiscent of the cryptococcal polysaccharide capsule, which is shed and causes immune modulation at a distance^50^ or can be retained around cells to play a fungal-protective role^51,52^. Based on these observations, we hypothesize that SOW could play a similar role during *Coccidioide*s infection. Clearly, much more study of SOW is called for, and it may be an excellent antifungal drug target that has yet to be exploited to treat coccidioidomycosis.

Finally, our study of the transcriptome of arthroconidia stocks generated and stored under various conditions indicates the wide variability of cell state for these spore forms. As arthroconidia are the starting material for most experiments in *Coccidioides*, these findings indicate the need to be cognizant of how experiments are designed and that the generation of arthroconidia is an important variable that needs to be controlled to attain reproducible results.

While is it likely not feasible to do all experiments with freshly-harvested arthroconidia, these data indicate that maintaining consistent duration of storage for comparable samples is important.

## Acknowledgements

This research was supported by the HHMI Hanna Gray Fellowship (to CH), the Program for Breakthrough Biomedical Research, which is partially funded by the Sandler Foundation (to CH), NIH R21AI172185 (to AS), and NIH U19AI166798 (to AS) for funding. AS is a Chan Zuckerberg Biohub – San Francisco Investigator. We also acknowledge the UCSF PCAT for use of equipment and UCSF CAT and CZI Biohub San Francisco for sequencing resources.

