## Supplementary figures and captions for "Optimizing in vitro spherulation cues in the fungal pathogen *Coccidioides*"

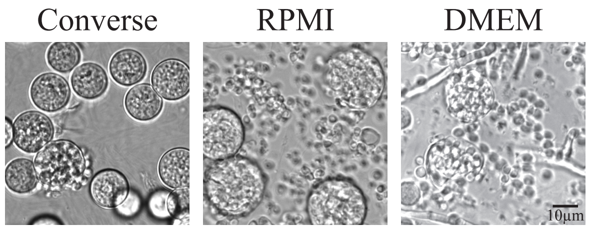


Figure S1

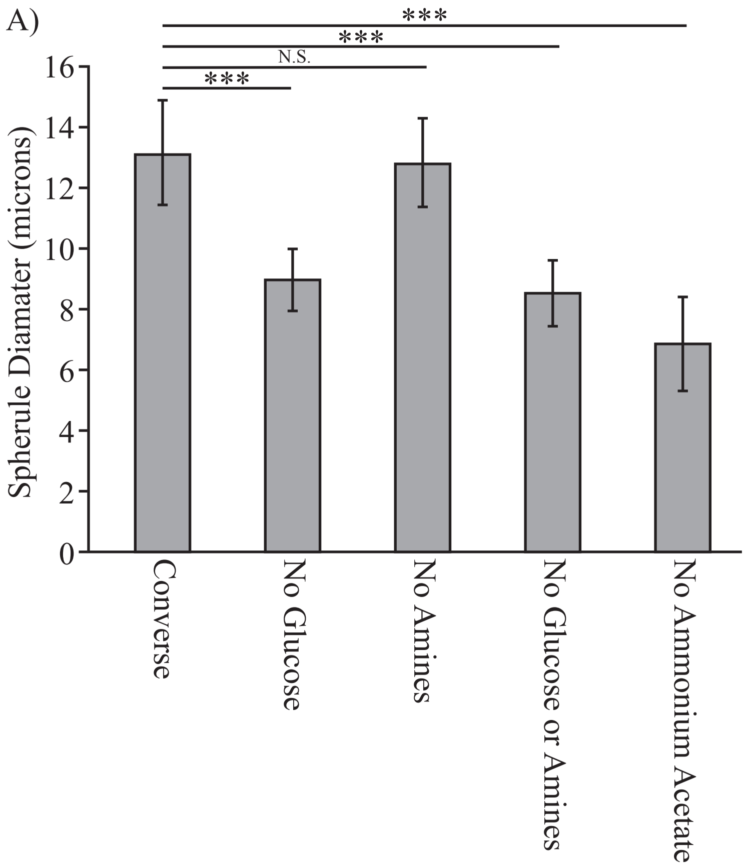

A)
***
***
N.S.
***
16
14
12
10
8
6
4
2
0
Spherule Diamater (microns)
Converse
No Glucose
No Amines
No Glucose or Amines
No Ammonium Acetate


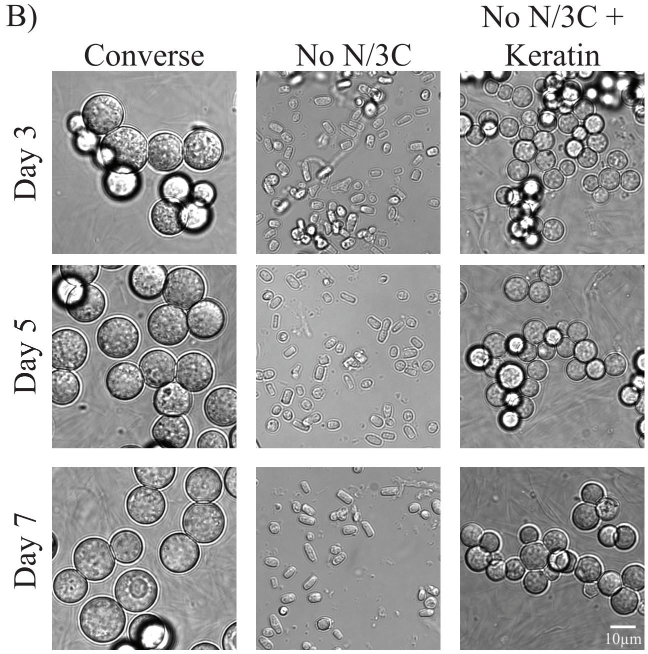

B)
Converse
No N/3C
No N/3C +
Keratin
Day 3
Day 5
Day 7
10μm


Figure S2

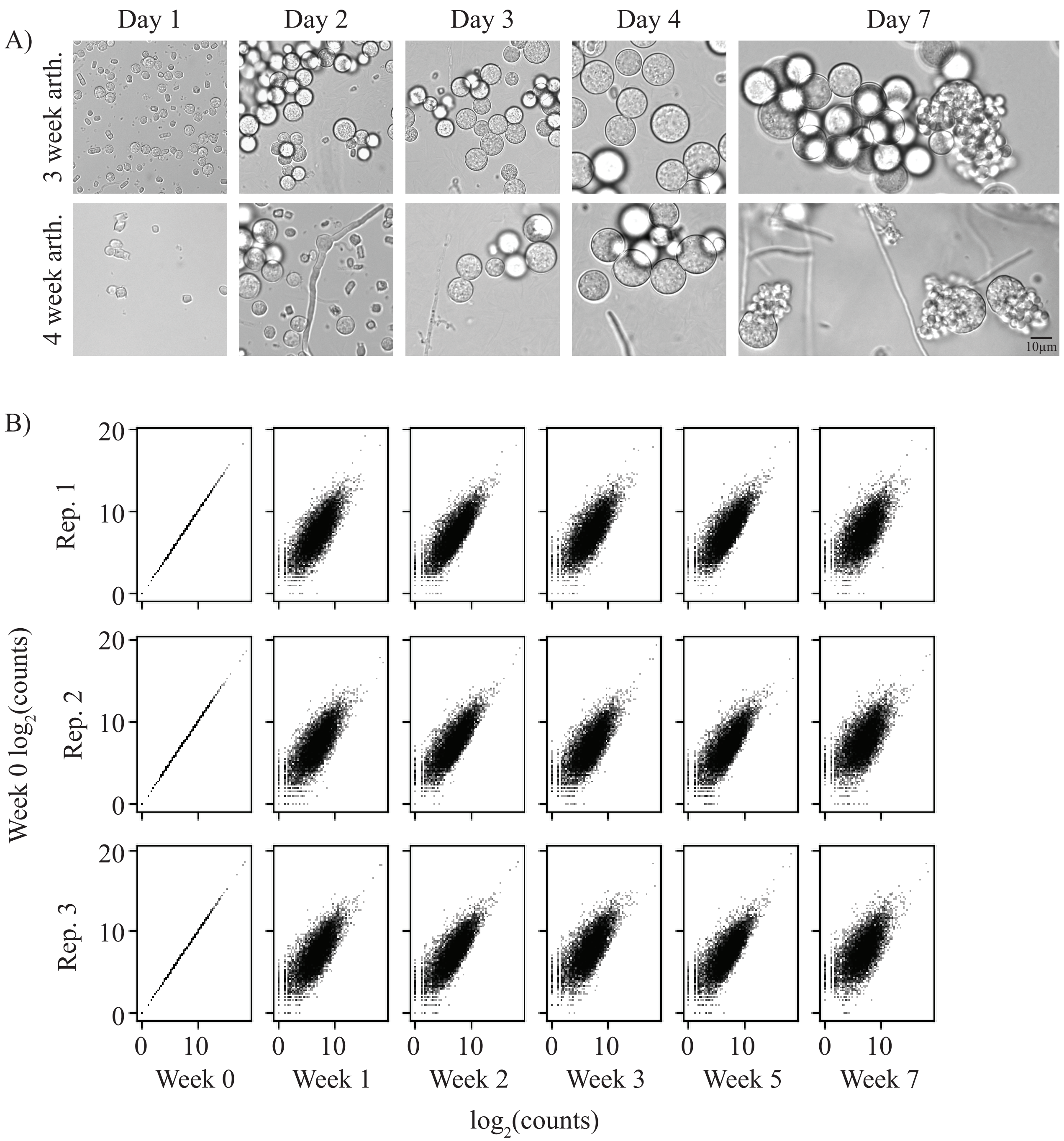
A)
Day 1
Day 2
Day 3
Day 4
Day 7
3 week arth.
4 week arth.
10μm
B)
Rep. 1
Rep. 2
Rep. 3
Week 0 log2(counts)
0
10
20
Week 0
Week 1
Week 2
Week 3
Week 5
Week 7
log2(counts)

Figure S3

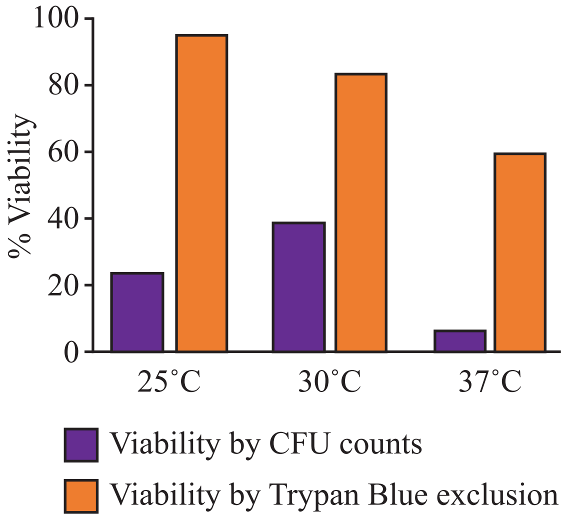
100
80
60
40
20
0
% Viability
25˚C
30˚C
37˚C
Viability by CFU counts
Viability by Trypan Blue exclusion

Figure S4

**Supplementary Material Captions**

**Figure S1: Spherulation in various media.** $10^6$ arthroconidia/mL were placed in Converse media + 0.5 % Tamol, RPMI + 10 % FBS, or DMEM + 20 % FBS under standard spherulation conditions (39˚C, 10% $CO_2$). Cells were fixed with 4 % PFA and images obtained on day 3 of incubation.

**Figure S2: Spherule comparisons for across different media types.** A. Average spherule diameter at day 7 for each condition in Figure 2A in microns. Spherule diameters were measured by hand in Fiji for at least 200 spherules per condition. *** $p < 0.0001$ by 2-sided t-test. B. $10^6$ arthroconidia/mL were grown in standard spherulation conditions (39˚C, 10 % $CO_2$) in either Converse or Converse media lacking ammonium acetate, glucose, N-Z amines with additional 3 % Keratin (No N/3C + Keratin) or without 3 % Keratin (No N/3C). Cells were fixed with 4 % PFA and monitored by light microscopy at day 3, 5, and 7 post-inoculation.

**Figure S3: Alterations in morphology and transcriptome caused by storage temperature**. A. Arthroconidia stored at 4˚C in PBS for 3 and 4 weeks were placed in standard spherulation conditions and fixed cells were monitored by light microscopy at day 1, 2, 3, 4, and 7 post-inoculation. Arthroconidia that had been stored for longer amounts of time exhibited more hyphal contamination of spherule cultures. B. Scatterplots of $\log_2$(counts) for each gene in freshly harvested Week 0 arthroconidia (y axis) versus arthroconidia stored for 1-7 weeks in PBS at 4˚C (x axis), demonstrating large-scale transcriptomic changes are not due to a normalization artifact.

**Figure S4: Viability assay demonstrates variability.** Percent of viable arthroconidia assessed by trypan blue exclusion (on day of arthroconidia harvest) or by CFU counts (after growth of arthroconidia stock for 72h) in independently-generated spore stock replicates from 6A. These replicate data demonstrate variability in this assay.

**Table S1: Tab-delimited text file containing transcript abundance ratios comparing arthro-conidia stored at 4˚C in PBS for multiple weeks**

Each row corresponds to a transcript. The columns are as follows: UNIQID: systemic gene name from Silveira genome[12]. Systematic gene names with _1, _2 appended have multiple isoforms as detected by kallisto although not all isoforms passed read count filter. NAME: Short gene name from Mandel et al[14]. Cp_anno: GenBank Cp Silveira annotation. CiRS: systematic CiRS gene name for the InParanoid-mapped *C. immitis* RS ortholog. CiRS_anno: Genbank annotation for CiRS ortholog. HcG217B: systematic HcG217B GSC gene name for the InParanoid-mapped *Histoplasma* G217B ortholog. HcG217B_anno: GSC annotation for HcG217B ortholog. The next 5 columns give limma adjusted p-values for differential expression for the listed contrasts. The next 18 columns give kallisto estimated normalized counts for each sample. The next 5 columns give limma generated $log_2$ fold change values for the listed contrasts.

**Table S2: Tab-delimited text file containing transcript abundance ratios comparing arthroconidia generated at 3 different temperatures**

Each row corresponds to a transcript. The columns are as follows: UNIQID: systemic gene name from Silveira genome[12]. Systematic gene names with _1, _2 appended have multiple isoforms as detected by kallisto although not all isoforms passed read count filter. NAME: Short gene name from Mandel et al[14]. Cp_anno: GenBank Cp Silveira annotation. CiRS: systematic CiRS gene name for the InParanoid-mapped *C. immitis* RS ortholog. CiRS_anno: Genbank annotation for CiRS ortholog. HcG217B: systematic HcG217B GSC gene name for the InParanoid-mapped *Histoplasma* G217B ortholog. HcG217B_anno: GSC annotation for HcG217B ortholog. The next 3 columns give limma adjusted p-values for differential expression for the listed contrasts. The next 18 columns give kallisto estimated normalized counts for each sample (6 replicates per temperature condition). The next 3 columns give limma generated $log_2$ fold change values for the listed contrasts.
